# Crossover Interference Mediates Multiscale Patterning Along Meiotic Chromosomes

**DOI:** 10.1101/2024.01.28.577645

**Authors:** Martin A. White, Beth Weiner, Lingluo Chu, Gyubum Lim, Nancy E. Kleckner

**Affiliations:** Department of Molecular and Cellular Biology, Harvard University, Cambridge MA, USA; Present address: Bioscience and Biomedical Engineering Thrust, Systems Hub, Hong Kong University of Science and Technology (Guangzhou), Guangzhou 511400, People’s Republic of China

## Abstract

The classical phenomenon of crossover interference is a one-dimensional spatial patterning process that produces evenly spaced crossovers during meiosis. Quantitative analysis of diagnostic molecules along budding yeast chromosomes reveals that this process also sets up a second, interdigitated pattern of related but longer periodicity, in a “two-tiered” patterning process. The second tier corresponds to a previously mysterious minority set of crossovers. Thus, *in toto*, the two tiers account for all detected crossover events. Both tiers of patterning set up spatially clustered assemblies of three types of molecules (“triads”) representing the three major components of meiotic chromosomes (crossover recombination complexes and chromosome axis and synaptonemal complex components), and give focal and domainal signals, respectively. Roles are suggested. All observed effects are economically and synthetically explained if crossover patterning is mediated by mechanical forces along prophase chromosomes. Intensity levels of domainal triad components are further modulated, dynamically, by the conserved protein remodeler Pch2/TRIP13.

## Introduction

Meiosis is the cellular program that gives haploid gametes from diploid progenitor cells as required for sexual reproduction. A core component of meiosis is the generation of genetic diversity. Shuffling of alleles between the maternal and paternal versions of each chromosome (“homologs”) is accomplished by crossover recombination^1^. As discovered over a century ago, crossovers occur at different positions in different nuclei but, nonetheless, tend to be evenly spaced along any given chromosome^2, 3^. This outcome implies that occurrence of a crossover at one position “interferes” with occurrence of another crossover nearby. This effect is evolutionarily important because even spacing, *per se*, enhances crossover-mediated allelic shuffling^4^. Crossover interference thus provides an interesting example of one-dimensional spatial patterning with significant evolutionary implications in a biological system.

Crossover patterning is imposed upon a large array of so-called “precursor” recombination complexes. These complexes comprise nascent interhomolog DNA interactions that link homolog structural axes all along the lengths of the chromosomes. The patterning process designates a subset of these precursors for eventual maturation into crossovers. It has long been assumed that a single basic process gives rise to a specific set of canonical “interfering crossovers” that are detectable as focal assemblies of crossover recombination complexes by electron or fluorescence microscopy (Fig. 1A). However, some organisms, including budding yeast, also exhibit a minority set of crossovers (as many as ∼30% of total crossovers in budding yeast) that are not marked by such focal assemblies. The existence of such events is revealed by genetic or DNA analysis, where the number of total detected crossovers is greater than the number of crossovers defined cytologically^5^. The basis for these minority events has been mysterious (below).

**Fig. 1.**
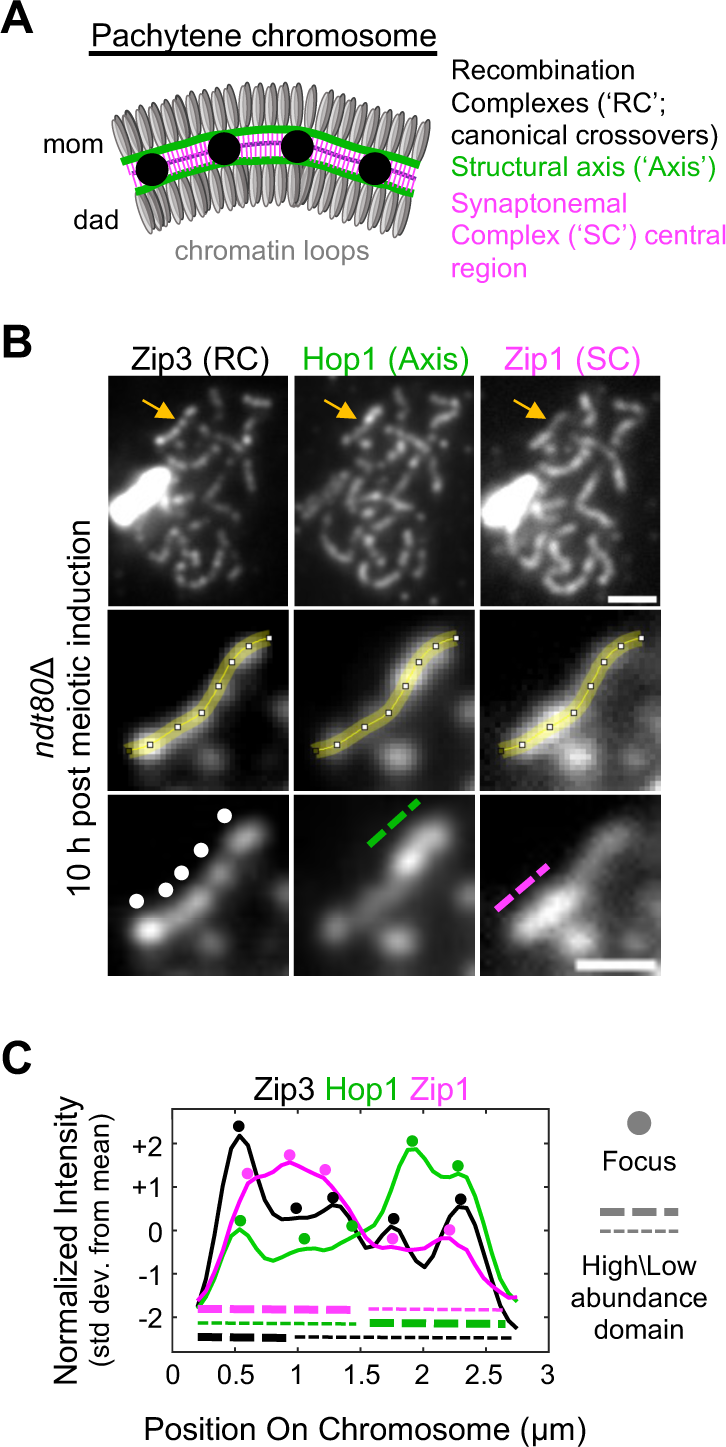
**Zip3, Hop1 and Zip1 signals are broad domainal fluctuations punctuated by focal peaks**. (**A**) Cartoon of pachytene chromosome structure, with homologs (‘mom’ and ‘dad’) and key components indicated. (**B**) *Top*, immunostained spread chromosomes from a pachytene arrested (*ndt80*Δ) cell. Arrows indicate example chromosome. Scale bar, 2 μm. *Middle*, example chromosome whose path was traced (yellow) to extract its intensity profiles. *Bottom*, example chromosome with Zip3 foci (dots) and Hop1 and Zip1 high abundance domains (dashed lines) indicated. Scale bars, 1 μm. (**C**) Normalized intensity profile of example chromosome. Std dev, standard deviations.

Spatial patterning of crossovers necessarily requires communication along chromosomes. All current models suggest that communication for crossover patterning involves transmission of a signal along underlying structures, either the axes of individual homologs and/or the synaptonemal complex (SC), a conserved structure that links homolog axes along their lengths^6^ (Fig. 1A). In budding yeast, canonical crossover interference is imposed prior to SC formation, with communication occurring along axes as implied by dependence on axis components Topoisomerase II and Red1^7, 8^. Communication along structures can confer the required effects because, universally, recombination complexes are physically and functionally associated with underlying chromosome structural elements at all stages^6^. The basis for this communication is unknown. One possibility is that it involves redistribution of mechanical stress along structural axes^9^ in a manner analogous to the patterning of inter-sister bridges along mitotic prophase chromosomes^6, 10^.

In the present study, we have further explored the relationship between chromosome structure and crossover localization by quantitatively defining and analyzing the spatial patterns of crossover-correlated recombination proteins and of axis and SC proteins along pachytene chromosomes of budding yeast. The observed effects have revealed that the basic classical crossover interference process sets up not one, but two tiers of patterns; that the two tiers together, account for both canonical and “minority” crossovers; and that both tiers set up “triads” comprising clusters of the three analyzed components. Chromosome structure is further altered by the conserved protein remodeler Pch2/TRIP13, which dynamically modulates the relative abundances of triad components. A synthetic model for two-tiered patterning and potential roles for the presence of triads are discussed.

## Results

### Recombination, axis, and SC components all exhibit both focal and domainal loading

Spatial patterns of crossover-correlated recombination proteins and of axis and SC proteins were examined along pachytene chromosomes of budding yeast (Fig. 1B top). Recombination proteins were represented by either Zip3 or Zip2, two molecules that specifically promote crossover formation and whose distribution of fluorescent foci define canonical crossover interference in this organism^5, 7^. Zip3 is a presumptive SUMO E3 ligase; Zip2 is a key member of a three-protein complex that directly mediates interaction between branched DNA molecules and chromosome structures^11^. Axes and SCs were represented, respectively, by the widely conserved HORMA domain protein Hop1^12^ and Zip1, yeast’s version of the universal SC central region transverse filament protein (Fig. 1A). Quantitative fluorescence intensity profiles for the three types of molecules were defined by tracing along the lengths of individual chromosomes (Fig. 1B middle, C).

Previous yeast studies have relied on contrast adjustment of fluorescence images to visually define peaks of Zip3 intensity (“foci”) as sites of evenly spaced crossovers^7^ and to define broader domainal distributions of Hop1 and Zip1 as presumptive manifestations of their general structural localization^13^ (e.g. Fig. 1B bottom). Quantitative tracing now reveals that all three molecules exhibit both features, with local peaks superimposed upon broader domains (Fig. 1C). Visual inspection of such traces gives the further impressions that: (a) foci and domains both seem to be evenly spaced; and (b) for both types of signals, the three molecules tend to colocalize with one another. These features, confirmed and extended by quantitative analysis below, capture the fundamental findings of this study.

### FFT analysis reveals two periodic patterns corresponding to ‘foci’ and ‘domains’

The intensities of Zip3, Hop1 and Zip1 signals were measured along 81 individual chromosomes of yeast meiotic cells arrested at the pachytene stage (using the *ndt80*Δ mutation). The component patterns present within these signals were revealed using Fast Fourier Transform (FFT) analysis. This method describes any given signal as the sum of multiple sine/cosine waves of different periodicities and amplitudes. The most prominent component periodicities in each trace appear as peaks in the corresponding plot of “power spectral density” (PSD) (Fig. 2A bottom).

**Fig. 2.**
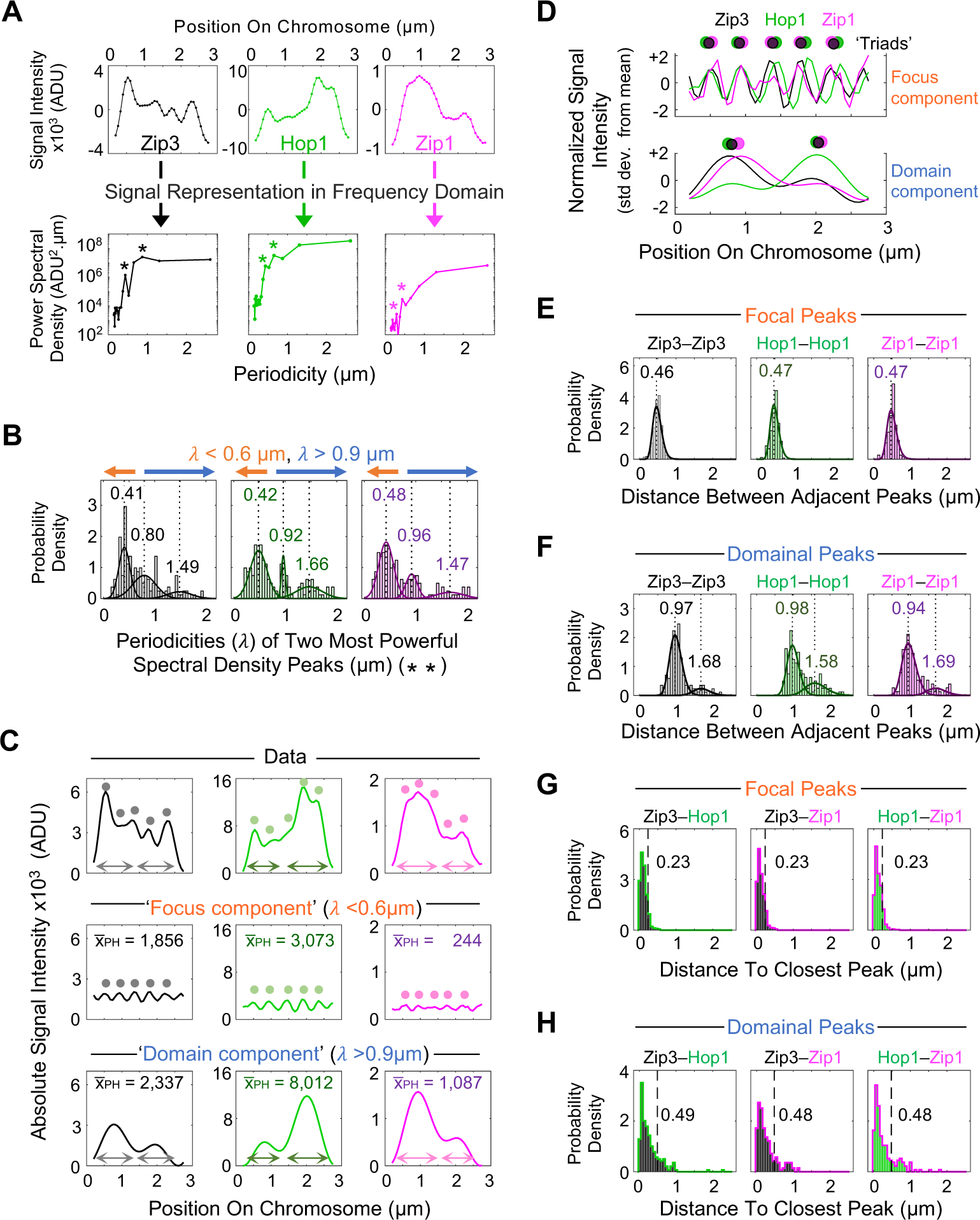
Frequency analysis reveals two component intensity periodicities corresponding to triads of focal and domainal signals. (**A**) Intensity profiles of example chromosome (same as Fig. 1B) and their corresponding power spectral density plots. Asterisks indicate two most prominent PSD peaks. ADU, analog digital units. (**B**) Distributions of two most powerful PSD peaks across all measured chromosomes (n = 81), with best fit 3-component Gaussian mixture models (solid lines) and their corresponding means (dashed lines) indicated. Orange and blue arrows indicate thresholds used for reconstruction of focal and domainal signals. (**C**) Intensity profiles, for the example chromosome (‘Data’) and its corresponding reconstructed focus and domain components along with their mean peak heights (xlJ_PH_). Dots, ‘foci’; arrows, ‘domains’. ADU, analog digital units. (**D**) Normalized intensity profiles of focus and domain component of example chromosome with Zip3/Hop1/Zip1 triads indicated. Std dev, standard deviations. (**E** and **F**) Distributions of distances between adjacent focal and domainal peaks, with best fit Gaussian distributions (solid lines) and their corresponding modes (dashed lines). (**G** and **H**) Distributions of distances between each Zip3 peak and its closest Hop1 peak (left) and analogous distributions for Zip3-Zip1 and Hop1-Zip1, for focal and domainal signal components. Dashed lines and corresponding values indicate expected medians if the two molecules were independently phased.

PSD plots were obtained for each intensity profile and the two most prominent PSD peaks in each plot were identified (e.g., Fig. 2A bottom, asterisks). Then, for each molecule, the distribution of the corresponding periodicities across the measured population of chromosomes was calculated (Fig. 2B). Three striking results emerge. (1) For each of the three molecules, the most prominent periodicities fall into three discrete groups. (2) The average periodicities of the three groups, as defined by the means of corresponding Gaussian distributions, are the same for each molecule (∼0.45 μm, ∼0.9 μm and ∼1.5 μm). (3) The intermediate periodicity is about twice the shortest periodicity, implying a mechanistic relationship. The longest periodicity is rare; its value is not statistically robust; and it likely reflects a failure to detect a few intermediate periodicity peaks (Supplementary Fig. 1A, B). The analysis below therefore considers the intermediate and longer periodicities as a single group.

The presence of shorter and longer periodicity groups implies the occurrence of two types of fluctuations in the corresponding signal intensities. These fluctuations can be visualized directly, for each molecule along each chromosome, by “inverse FFT analysis”. For this purpose, FFT periodicities shorter than 0.6 μm, or longer than 0.9 μm, were selected (omitting periodicities that might belong to either group) (Fig. 2B, orange and blue arrows). The intensity profiles resulting from the fluctuations in each group along each individual chromosome were then reconstructed (representative examples in Fig. 2C). These reconstructions reveal that the shorter periodicity fluctuations correspond to the visually identifiable focal component while the longer periodicity fluctuations correspond to the visually identifiable domainal component (Fig. 2C).

This analysis also reveals that, for all three molecules: (i) focal peaks are narrower than domainal peaks and (ii) focal and domainal peaks fluctuate around lower and higher mean values, respectively (Fig. 2C, ‘xlJ_PH_’). Interestingly, focal peaks of Zip3 are relatively more prominent relative to their domainal peaks than are peak of Hop1, which in turn are relatively more prominent than those of Zip1. This hierarchy explains why, in previous studies, which involved visual thresholding of images, Zip3 intensities were defined as foci; Hop1 intensities were defined as a combination of foci and broad domains; and Zip1 intensities were defined as broad domains^13, 14^.

### Focal and domainal signals are evenly spaced triads, each comprising closely spaced assemblies of crossover recombination, axis and synaptonemal complex components

Reconstructed intensity profiles support the impression (Fig. 1C) that peaks of all three molecules colocalize for both focal and domainal signals, which thus comprise two evenly spaced arrays of triads. This is clear by visual inspection of inverse FFT outputs for all 81 examples of each type (e.g., Fig. 2D and Supplementary Fig. 1C). This conclusion is further supported by three lines of evidence.

- Distances between adjacent peaks of the same molecule show the same modal distance for all three molecules (∼0.45 μm and ∼0.9 μm for focal and domainal signals respectively; Fig. 2E, F). Also, peaks of both types are similarly evenly spaced as defined by best fit gamma distribution shape parameters (Supplementary Fig. 1D).
- If peaks of different molecules are clustered in triads, a peak of a given molecule and the nearest peak of a different molecule should occur within a triad and thus be closer together than adjacent peaks of the same molecule. Accordingly, the modal distance between nearest heterologous peaks is much lower (67 nm; 1 pixel) for both focal and domainal peaks (Fig. 2G, H) versus the distances between nearest peaks of the same molecule (∼0.45 μm and ∼0.9 μm; Fig. 2E, F). The modal distance of 67 nm for nearest peaks of different molecules implies very tight clustering of the three component peaks within each triad.
- In principle, two evenly spaced arrays of peaks might be phased coordinately (e.g. with the three types of peaks clustered into triads) or independently. In the former case, the distribution of distances between each peak of one type and the nearest peak of a second type should be nonuniform. In the latter case, such distributions should be uniform, ranging from 0 to ∼0.25 μm for shorter periodicity signals and 0 to ∼0.5 µm for longer periodicity signals. We find that all such distributions are nonuniform (Fig. 2G, H and Supplementary Fig. 1E, F).

All the above features were confirmed in an analogous experiment using Zip2 rather than Zip3 as a marker for recombination complexes and with pachytene nuclei obtained at an appropriate time point in wild type (*NDT80*) cells undergoing synchronous meiosis, rather than in *ndt80*Δ arrest conditions (Supplementary Fig. 2).

### Focal and domainal triad patterns are both downstream outcomes of crossover interference

Crossover interference is defined by Coefficient of Coincidence (CoC) analysis^2, 3^. Chromosomes are divided into intervals which are then examined pairwise for the frequency of chromosomes that exhibit two crossovers versus the frequency expected on the basis of independent occurrence^3, 15^. The diagnostic effect of interference is that double crossovers are much rarer than expected at smaller inter-interval distances (CoC <<1, and zero at very short distances) and rise to the level expected for independent occurrence (CoC ∼1) over some characteristic longer distance. Both focal and domainal triads exhibit interference, as seen by analysis of reconstructed patterns for all three individual components. Domainal peaks show stronger interference than focal peaks, as manifested in the inter-interval distances at which CoC rises to 0.5^16^ (L_coc_ = ∼0.65 μm and ∼0.3 μm, respectively) in accord with their longer periodicity (Fig. 3A). Identical results are obtained for Zip2 foci in the absence of pachytene arrest (Supplementary Fig. 3A, B).

**Fig. 3.**
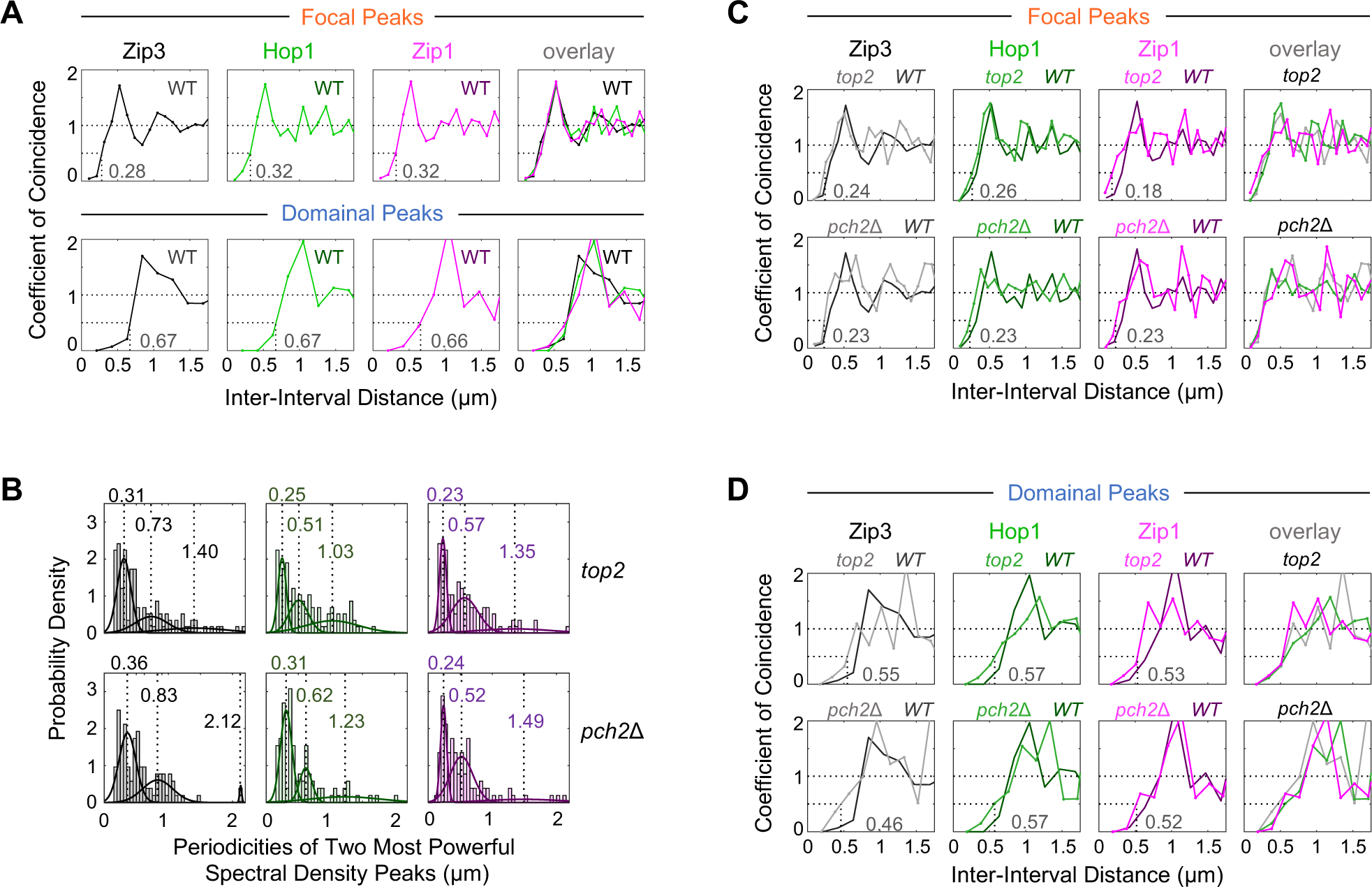
Crossover interference governs the patterns of both focal and domain triads. (**A**) Coefficient of coincidence (CoC) curves for focal and domainal peaks of chromosomes isolated from an *ndt80*Δ strain (‘WT’) with corresponding L_coc_ values. (**B**) Distributions of the two most powerful PSD peaks across all measured *pCLB2-TOP2* (‘*top2*’, n = 60) and *pch2*Δ (n = 52) chromosomes, with best fit 3-component Gaussian mixture models (solid lines) and their corresponding means (dashed lines). CoC curves for focal (**C**) and domainal (**D**) peaks of chromosomes isolated from *top2* and *pch2*Δ strains with corresponding WT CoC curves (**A**) for reference.

In budding yeast, canonical interference is reduced by depletion of Topoisomerase II (‘Top2’; *pCLB2-TOP2*^7^). Interference is also reduced in some situations by elimination of the conserved protein remodeler AAA+-ATPase Pch2/TRIP13^14, 17^ (*pch2*Δ). We now find that, in both mutant conditions (n = 60 and 52 pachytene chromosomes, respectively), focal and domainal triad patterns still occur, just as in wild type, by all criteria (Fig. 3B and Supplementary Fig. 3C – F). However, also in both conditions, the strength of interference is reduced, for both sets of triads, as shown by shifting of corresponding CoC curves to smaller inter-interval distances (Fig. 3C, D). Correspondingly: (i) the average periodicities of all PSD groups are smaller than those of wild type (compare Figs. 3B and 2B); and (ii) all average homologous inter-peak distances are reduced, as expected if double crossovers tend to be closer together (Supplementary Fig. 3G).

### Focal and domainal triads correspond to canonical and minority crossovers, respectively, thereby defining the sites of all crossovers in wild type meiosis

#### Focal triads correspond to canonical crossovers

Focal Zip2 and Zip3 peaks, as defined by FFT reconstruction analysis, exhibit the same spacing and CoC relationships as those for canonical crossover interference in published data (Fig. 4A, B). These focal peaks occur in triads that also contain Hop1 and Zip1, with all component molecules showing the same spacings and CoC relationships as one another (above). Furthermore, the magnitude of the shift in the CoC curves seen for focal triad components in *pCLB2-TOP2* (Fig. 3C) is the same as that described previously for canonical interference of Zip3 foci (Supplementary Fig. 4A). Thus, focal triads occur at sites of canonical interfering crossovers.

**Fig. 4.**
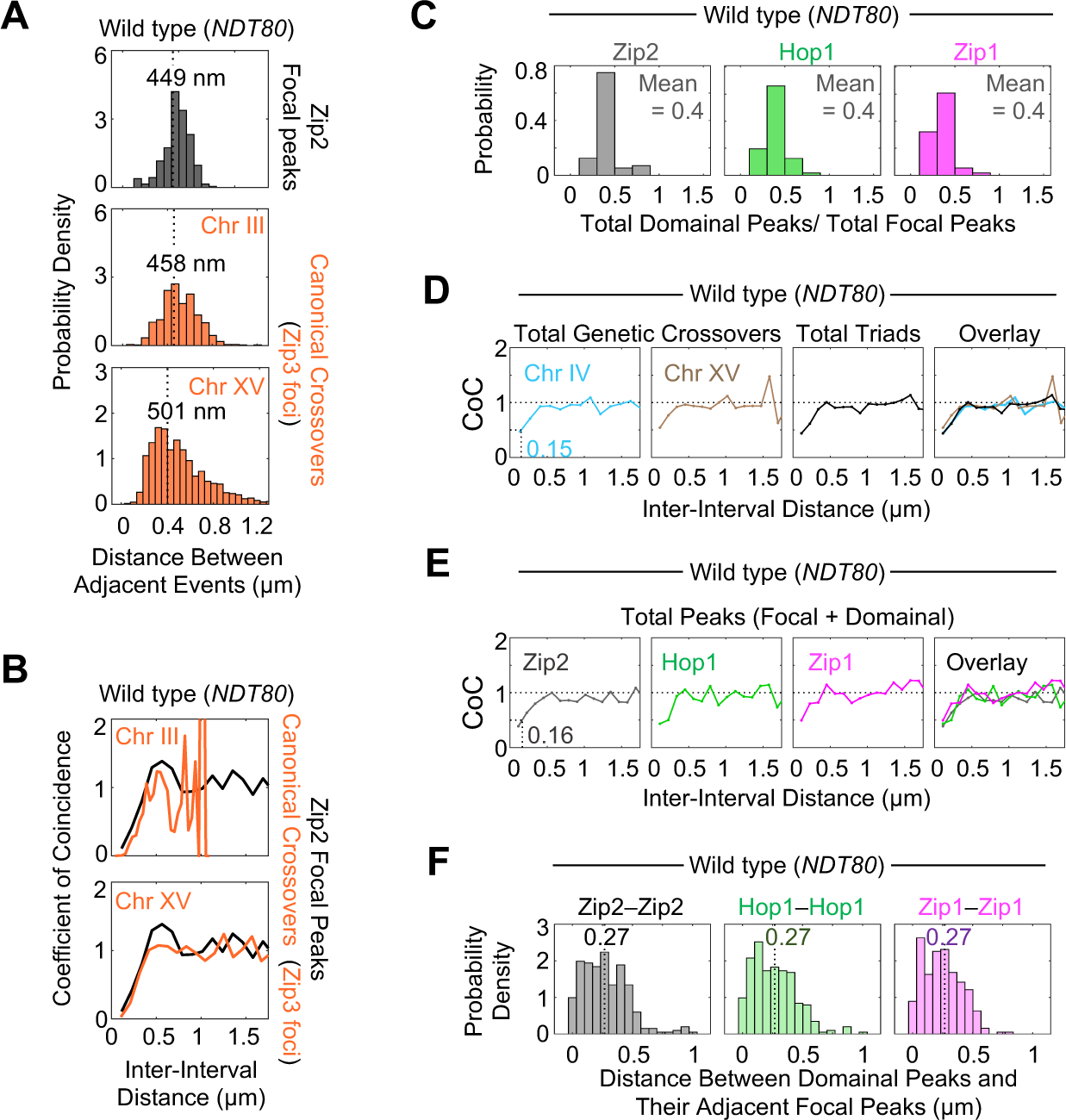
Focal triads correspond to canonical crossovers and domainal triads correspond to minority crossovers. (**A** and **B**) Comparison of Zip2 focal peaks (n = 56 chromosomes; this study) and published canonical crossovers^7^ (n = 683 and 822 for chromosome (Chr) III and XV) in a wild type (*NDT80*) genetic background. (**A**) Distributions of distances between adjacent events with modes of best fit gamma distributions indicated. (**B**) Coefficient of coincidence (CoC) curves. (**C**) Distribution of ratios of total domainal to total focal peaks for each molecular signal. (**D**) Comparison of CoC curves for published total genetic crossovers^16, 18, 19^ (n = 72 for Chr IV and Chr XV) and total triads (this work; total focal and domainal peaks; average for Zip2, Hop1 and Zip1 signals). For genetic crossovers, distances in DNA bp were converted to μm chromosome length assuming 324 bp/nm^16^. (**E**) CoC curves for all peaks (focal and domanial) along measured chromosomes, with corresponding L_coc_ value. (**F**) Distribution of distances between domainal peaks and their adjacent focal peaks with medians (dashed lines) indicated.

#### Domainal triads correspond to minority crossovers

Wild type meiosis exhibits both majority canonical crossovers and a minority residuum of “non-canonical” crossovers (above).

Three lines of evidence suggests that domainal triads occur at the sites of these minority non-canonical crossovers.

- First, in yeast, the ratio of the two types of crossovers is 0.4:1.0^16, 18, 19^. Exactly this same ratio is observed for longer periodicity triads versus shorter periodicity triads (mean ratios of 0.4 for all components; Fig. 4C and Supplementary Fig. 4B).
- Second, the CoC curves for total crossovers and for total triads are strikingly similar. The CoC curve for total crossovers as defined by DNA sequence analysis along budding yeast chromosomes IV and XV^16, 18, 19^ (Fig. 4D left panels) differs significantly from that of canonical, cytologically detected crossovers (also defined for chromosome XV (Fig. 4B)), in two diagnostic respects. (a) The strength of interference is much weaker for total events than for canonical events alone, with a substantial shift of the CoC curve towards smaller inter-interval distances. (b) For total crossovers, the CoC curve does not fall to a value of zero at the smallest inter-interval distances. The CoC curves for total triads (focal + domainal) are virtually identical to those for total DNA-defined crossovers in both respects, as defined for each individual component and their average (Fig. 4D, E and Supplementary Fig. 4C).
- Third, the failure of a CoC curve to fall to zero implies that detected crossovers sometimes occur very close together along the same chromosome. Correspondingly, the adjacent focal and domainal peaks of any single component are often very close together, with half of all adjacent pairs separated by less than 0.27 μm (Fig. 4F). This relationship is explained by the fact that crossovers of the two types can occur at adjacent precursor sites, which occur at the same average distance (∼233 nm; Supplementary Note).

#### Implications

Given that focal and domainal peaks correspond to canonical and minority crossover sites, the two sets of triads account for all the crossovers that occur in wild type meiosis. Moreover, both types of crossovers are downstream of interference, thus invalidating the previous speculation that minority crossovers are “non-interfering (Supplementary Discussion). Furthermore, Zip2 and Zip3 (and also Zip1) are members of a set of molecules known to promote progression of canonical crossovers (“ZMMs”). Since these molecules are present, analogously, in both focal and domainal triads, ZMMs may be required for formation of both types of crossovers. Indeed, in mouse, which also has both majority and minority crossovers, all crossovers require the ZMM protein MutSγ^20^.

### Pch2 dynamically modulates triad component levels throughout pachytene

Previous work reported that the relative intensities of Hop1 and Zip1 tend to differ along the lengths of the chromosomes over long length scales^13^ (e.g. Fig. 1B) and that absence of conserved meiotic protein remodeler Pch2/TRIP13 abrogates this tendency, with the two proteins now exhibiting a strong tendency to occur at similar, and elevated, levels all along the chromosomes^13, 21^ (e.g., Fig. 5A, B, C left). These effects can be quantified by measuring the relative levels of each pair of molecules, defined as the ratio of the normalized intensities of Hop1 and Zip1 along each chromosome (Fig. 5C right). Greater and lesser fluctuations in wild type and *pch2*Δ are revealed by the extent to which this ratio deviates from the mean value (i.e. the standard deviation, ‘SD’; e.g. for a single chromosome in Fig. 5C right and for all measured chromosomes in Fig. 5D).

**Fig. 5.**
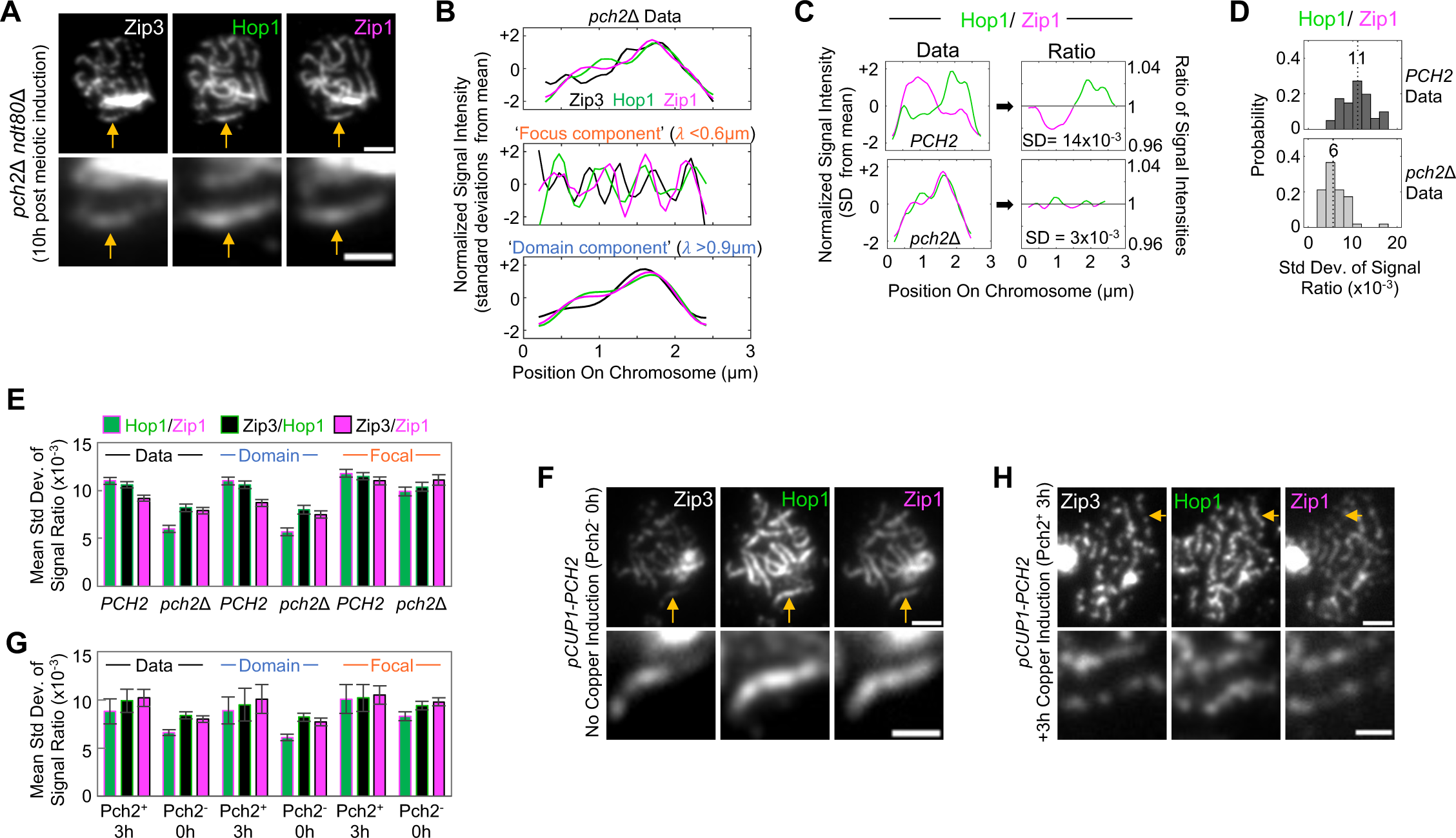
**Pch2 modulates the relative abundances of triad components**. (**A**, **F**, and **H**) Immunostained spread chromosomes (*top*; scale bar 2 μm) and example chromosome indicated by arrow (enlarged, *bottom*; scale bar 1 μm) from a pachytene arrested (*ndt80*Δ) *pch2*Δ cell (**A**) and inducible Pch2 cells grown in either the absence of Pch2 expression (**F**), or with 3h of Pch2 expression at pachytene (**H**). (**B**) Normalized intensity profiles of example *pch2*Δ chromosome (**A**, *bottom*). (**C**) Hop1 and Zip1 intensity profiles of example *PCH2* and *pch2*Δ chromosomes, with their corresponding signal ratio plots and standard deviation (SD) of signal ratio. (**D**) Distribution of standard deviations of Hop1/Zip1 signal ratios, with medians (dashed lines) indicated. Std Dev., standard deviation. (**E** and **G**) Mean standard deviation of signal ratio for each pairwise comparison of triad components. Error bars, standard error of the mean; n = 81, 52, 8, and 61 for *PCH2*, *pch2*Δ, Pch2^+^ 3h, and Pch2^-^ 0h respectively.

Corresponding analysis for all three pairs of signals in focal and domainal triads reveals that in wild type meiosis, this tendency for domainal variation is observed for all three molecules in all pairwise combinations and is reflected primarily in intensity differences within domainal triads (Fig. 5E (*PCH2*), 1C and Supplementary Fig. 1C). This matches the fact that these triads have the appropriate length scale and intensity differences (e.g. Fig. 2C). Furthermore, since absence of Pch2 does not alter the occurrence of triad structure (above), it must exert its effects by modulating molecular intensities within individual triads. Specifically, absence of Pch2 exerts its effects primarily on relative intensities within domainal triads, with lesser effects on focal intensities (Fig. 5E), in accord with the patterns observed in wild type. Interestingly, the effects of Pch2 are greatest for Hop1/Zip1, intermediate for Zip3/Hop1 and least dramatic for Zip3/Zip1 (Fig. 5E).

We further find that induction of Pch2 in a pachytene *pch2*Δ cell can restore the normal pattern of alternating Hop1/Zip1/Zip3 domainal hyperabundance dynamically in real time. In a *pch2*Δ *ndt80*Δ strain carrying a copper-inducible Pch2 expression construct (*pCUP1-PCH2*), arrested at pachytene in the absence of copper, chromosomes exhibit the typical *pch2*Δ phenotypes (n = 61 pachytene chromosomes; Fig. 5F, G ‘Pch2^-^ 0h’ versus Fig. 5E ‘*pch2*Δ’). If Pch2 expression was then induced and cells incubated for a further 3 hours, chromosomes return to a typical *PCH2* phenotype (n = 8 pachytene chromosomes; Fig. 5H, G ‘Pch2^+^ 3h’ versus Fig. 5E ‘*PCH2*’). We infer that, in wild type meiosis, Pch2 modulates the levels of triad components continuously throughout pachytene without affecting the existence of basic triad structure *per se*.

### Two-tiered patterning and correlated intensity distributions are synthetically explained by the “beam/film” stress hypothesis for crossover interference

We previously described a model in which crossover patterning is driven by mechanical effects along a linear bi-layered system^6, 9, 15, 16^. The original model considered a physical analogy in which a metal beam was coated by a thin brittle film with flaws. Stress would arise along the interface between the two layers due to differential expansion of the beam (e.g. in response to heating). A first crack would occur, perpendicular to the beam, triggered by stress at the weakest flaw site. The immediate effect is necessarily a local reduction of stress. That effect then intrinsically redistributes outward, dissipating with distance, giving a zone of reduced stress. Any subsequent (stress-promoted) cracks will tend to occur outside this zone of stress relief, ultimately giving an array of evenly spaced cracks separated by a characteristic “stress relief distance” (Fig. 6A left-middle).

**Fig. 6.**
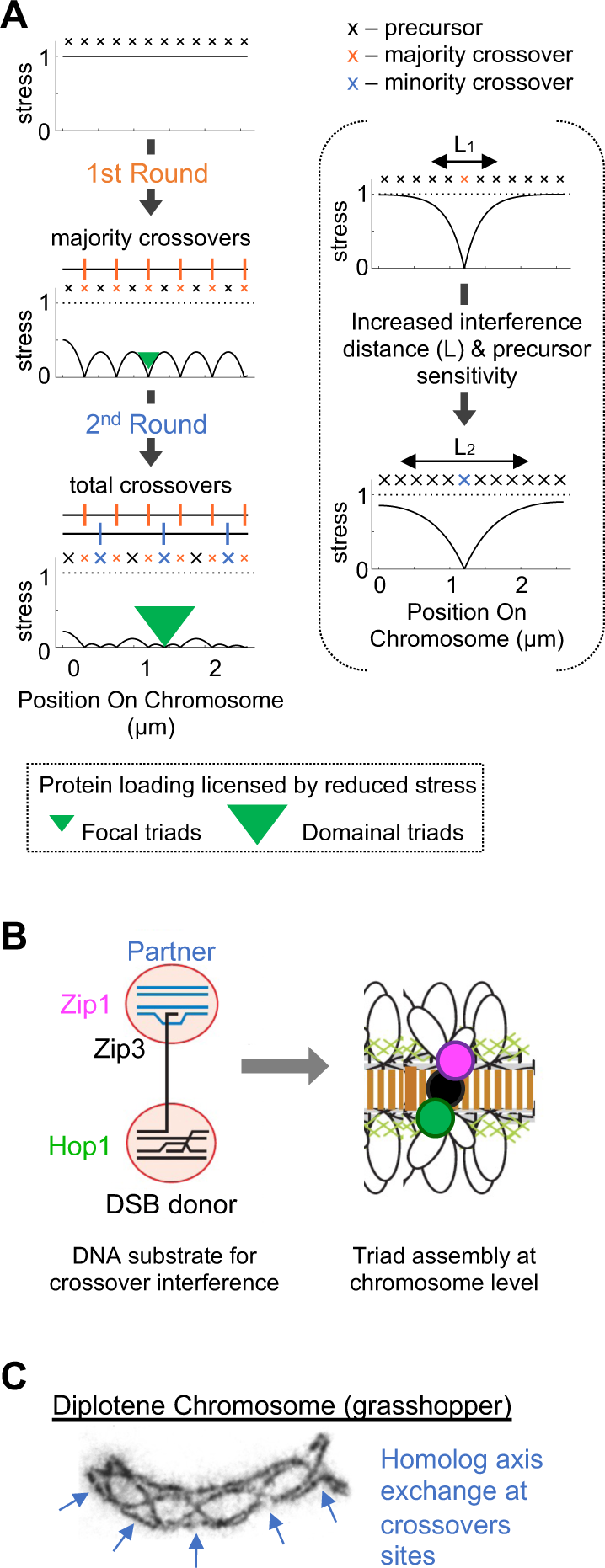
Two rounds of interference and roles of triads. (**A**) Two tiers of crossover interference are explained by two rounds of stress-promoted patterning in a bilayered system with flaws (text). (**B**) Model for how triads may regulate DNA events involved with crossover recombination. DSB, DNA double-strand break. (**C**) Electron micrograph of grasshopper late meiotic prophase (diplotene) chromosome^43^ with interhomolog axis exchanges indicated.

We propose that chromosomes form an analogous system^6, 9^. The chromatin that emanates from axes as loops (Fig. 1A) would act as the beam; the chromosome axis, which is an elastic meshwork of protein/protein/DNA/DNA interactions, would act as the film; and axis-associated “precursor” recombination complexes would behave as flaws. Pushing forces among adjacent chromatin loops would create mechanical stress along the axes. And the event promoted by this stress would be a molecular change that commits the affected recombination precursor to becoming a crossover. A closely related scenario can explain emergence of evenly spaced bridges between sister chromatid axes along mitotic post-prophase chromosomes, where morphological features point strongly to the presence of stress along chromosome axes^6, 10, 22^.

The existence of two tiers of crossover patterning is economically explained by two successive rounds of events that occur according to the “beam/film” scenario (Fig. 6A left). Simulations based on this scenario accurately recapitulate experimental CoC curves (Supplementary Fig. 5A, B). This model requires that two changes occur between the successive rounds (Fig. 6A right): (1) chromosomes must become stiffer for stress relief at minority crossover designations (i.e. interference) to propagate over a larger distance than during the first round; (2) unreacted precursors must become more sensitive to stress to enable their differential response at this second stage. These two requirements can be explained by formation of the synaptonemal complex (SC). In yeast, SC installation is nucleated at canonical crossover designation sites^6^ and thus will be a direct outcome of the first round of patterning, thus setting the stage for a second round. The SC adds crosslinks between homolog axes (Fig. 1A), which is predicted to increase their stiffness. Increased precursor sensitivity might be increased, for example, by association of previously axis-associated recombination complexes now with the SC.

This scenario also has the attractive feature that it could directly explain why canonical crossovers manifest as narrow ‘foci’ whereas minority crossovers manifest as broad ‘domains’. If loading of triad components is inhibited by stress and, thus, licensed by stress relief, their chromosomal loading after each round of patterning will mirror the distance and magnitude of the two successive resultant stress distributions (Fig. 6A left; green arrowheads). The relative magnitudes of components within triads would then be further modulated by Pch2/TRIP13 (above).

## Discussion

Crossover interference has long been assumed to generate a specific set of “interfering crossovers” that are evenly spaced along the chromosomes. Here, quantitative analysis of the intensity distributions of relevant molecules reveals that the basic interference process actually sets up two distinct tiers of spatial patterning which, together, account for all detected crossovers.

This finding emerges from the observation that the three basic types of chromosome structure components (recombination-related and axis and synaptonemal complex-related) occur in clustered local assemblies (“triads”) and that these triads are of two types: focal assemblies that occur with ∼450 nm spacing and define the patterning tier corresponding to canonical crossovers, and broader domainal assemblies that occur with ∼900 nm spacing and define the patterning tier corresponding to sites of previously-mysterious “minority” crossovers.

We note that while the latter set of crossovers was previously assumed to lack interference, we find instead that they do display interference, and over a greater distance than canonical crossovers. The previous view was based on modeling studies, which can mimic certain features of total crossover distributions by sprinkling some non-interfering crossovers randomly among canonical crossovers (Supplementary Discussion). The situation revealed here for yeast could apply to other organisms. In *Arabidopsis thaliana,* maize, mouse, humans, and possibly also tomato, wild type crossover patterns are well explained by the existence of two classes of crossovers that differ in interference behavior^23, 24, 25, 26, 27^ and can be explained analogously to the situation in yeast described here (Supplementary Discussion). Also, in mouse, all crossovers require the key “ZMM” protein MutSγ^20^, in accord with the hint that Zip3 may be involved in both majority and minority crossover sites in yeast (above).

### Mechanism of two-tiered crossover interference

Current models for crossover interference fall into two categories according to whether the communication required to set up the relevant pattern is mediated by redistribution of mechanical stress or by biochemical diffusion, notably by the process of Ostwald ripening of meiosis-specific E3 ligases such as Zip3^6, 28^.

The possibility of a mechanical model is directly supported by studies of mitotic prophase chromosomes, whose organization is analogous to that of meiotic prophase chromosomes, and which show clear evidence of mechanical stress along their axes which promotes morphological changes with appropriate hallmarks^10, 22^. We describe above a specific version of the previously described “beam-film” model which draws on this analogy to account for meiotic crossover patterning. We also show that two-tiered patterning of triads can be economically explained by this model (Fig. 6A and Supplementary Fig. 5A, B). In this context, a first round of patterning gives canonical ∼450 nm crossover sites; a change in chromosome state then occurs which allows a second round of patterning in which stress redistributes over longer distance, giving ∼900 nm minority crossover sites. This change in state could be installation of the synaptonemal complex which is a tightly coupled downstream outcome of canonical crossover patterning in budding yeast. The proposed model also has the attractive feature that it can directly explain why canonical crossovers manifest as narrow ‘foci’ whereas minority crossovers manifest as broad ‘domains’. In brief, the two types of intensity patterns would be dictated by the residual stress distributions after each tier of patterning. In summary, the stress hypothesis provides a synthetic and economical explanation for both the spacing patterns and intensity distributions for two-tiered crossover site triad formation.

In contrast, a diffusion-based mechanism might not yield the observed outcome so easily. For example, a prominent feature of two-tiered patterning is that events from the two tiers are interdigitated but do not directly interact/interfere with one another. In our proposed model, this effect is an automatic, built-in consequence of the fact that changes in stress can propagate along the “beam” (the chromatin) regardless of whether it does or does not encounter a “crack” (crossover designation) within the “film” (the axis). In a diffusion-based model, during a second round of patterning, the diffusing entity would need to be able to skip over/ignore designation sites created by the first round. This feature would require the additional assumption of some type of molecular differentiation between the two rounds. We also note that mechanical effects could define sites of crossovers and that diffusion-mediated condensation^6^ might then result in accumulation of specific molecules (such as E3-ligase Zip3) at those designated sites.

We further note that a key advantage of stress-mediated patterning is that the interference distance can be tuned by the stiffness of the chromosome feature along which interference propagates (e.g. the chromosome axis) ^9^. This feature underlies our proposition that an increase in chromosome axis stiffness is responsible for the increased interference distance between the two patterning tiers. In the case of minority crossovers, this increase might reflect increased stiffness due to SC formation (above). However, differences in stiffness could analogously explain differences in interference distances for canonical crossovers in diverse organisms across evolution. In this more fundamental case, stiffness could be modulated by effects on the chromosome axis. For this complex meshwork of protein/protein/DNA/DNA interactions, any increase in crosslinks could achieve the desired effect.

### Roles of triads

It is striking that both types of crossover sites exhibit assemblies of the three major types of meiotic chromosome components. We can propose two not-mutually-exclusive possibilities for the nature and role(s) of these triads.

- Model 1 (Fig. 6B): Meiotic recombination initiates by programmed DNA double-strand breaks (DSBs). The substrate for crossover designation is thought to comprise one (leading) DSB end in a nascent D-loop complex that is associated with the corresponding homolog partner axes while the other (lagging) DSB end remains associated with the axis of the originating homolog. At sites of canonical crossovers, crossover designation is likely imposed on the leading DSB end and, concomitantly, nucleates SC formation. The lagging end is brought into the developing complex only later, in “second end capture”. We suggest that crossover designation triggers a Zip2/3 assembly on the recombination complex *per se*; a Zip1 assembly on the leading end to stabilize interaction with the SC; and a Hop1 assembly on the lagging end to mediate regulated release for second end capture, in accord with Hop1’s intimate relationship to axis-associated cohesin complexes^12, 29^. Similar roles could pertain at minority crossover sites.
- Model 2: Crossing-over requires not only reciprocal exchange between homolog chromatids at the DNA level but also creation of continuous chromosome structure at the corresponding positions (“axis exchange”)^6^, Fig. 6C. Crossover-specific recombination complexes and bits of SC remain colocalized at crossover sites during this process, with SC known to play a significant role^30^. Hop1 localization to these sites has not been examined, but might be suspected, where it could mediate separation of sister chromatid axes and/or promote development of new axis structure at these sites.

### Modulation of molecular intensities by Pch2

Pch2 is not involved in setting up triad patterns or triad structure. It is only involved in modulating the levels of molecular components within triads. These molecules undergo several direct and indirect interactions which make it difficult to propose a specific mechanism. For example, yeast Pch2 can act directly on Hop1 to promote both its loading onto and removal from chromosomes^12, 31^. However, Pch2 is only seen loaded onto chromosomes in the presence of Zip1^32, 33^ and its absence is known to increase the levels of both Hop1 and Zip1^21^. Furthermore, Zip1 is known to promote re-localization of Zip3^34^.

Interestingly, however, in *pch2*Δ, all three molecules occur at the same relative levels in all triads all along the chromosomes, in both types of triads (Fig. 5B). In the presence of Pch2, the same is true for focal triads while, in domainal triads, different components occur at different relative levels. These effects suggest that there is a Pch2-independent default option in which the levels of all components are coupled such that they always occur in the same relative levels. Pch2 would then abrogate this coupling, differentially in domainal triads. A further interesting finding from the current work is that Pch2 can act dynamically to restore differential intensity levels even after the Pch2-independent condition has been established. We infer that such effects operate during development of pachytene chromosomal intensity patterns in wild type meiosis. Correspondingly, we can speculate as to how dynamic effects might play a key role in creating the observed effects (Supplementary Fig. 6A, B). We note that previous work involving a bulk-population assay (chromatin immunoprecipitation) revealed that Hop1 tends to be enriched at chromosome ends in a Pch2-dependent manner^35^. Our data show that Hop1 distributions vary considerably along both wild type and *pch2*Δ chromosomes without any reproducible pattern of terminal enrichment at the per-chromosome level (Supplementary Fig. 6C). However, a population-level tendency can be seen when data for all chromosomes is averaged, and we can further suggest that this effect is attributable primarily to the dispositions of domainal triads (Supplementary Fig. 6D).

## Materials and Methods

### S. cerevisiae strains

All strains used in this study are *MATa/MATα* derivatives of SK1.

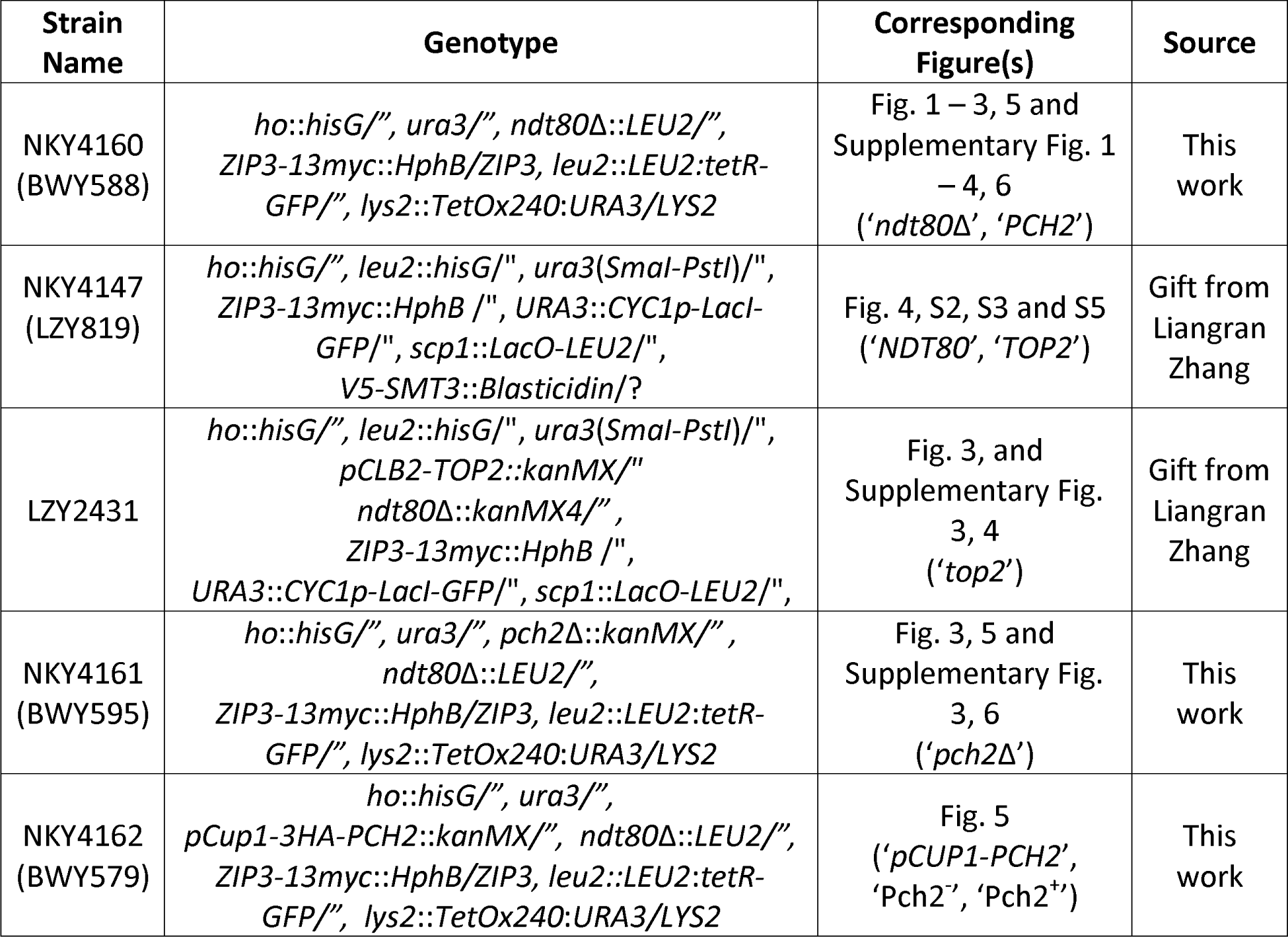

### Meiotic Induction

Mitotically growing cells were induced to undergo synchronous meiosis at 30 °C using the previously described SPS method^36^. Time 0 was defined as the transfer of cells to sporulation medium. For *NDT80* strain LZY819, cells were harvested for chromosome spreading 6 h later, when pachytene cells were most abundant. For *ndt80*Δ strains BWY588, LZY2431, and BWY595; cells were harvested 10 h post meiotic induction for maximal accumulation of pachytene cells. For *pCUP-PCH2 ndt80*Δ strain BWY579, a sample of cells (‘No Copper induction’) was first harvested 10 h post meiotic induction. Copper was then added to the growth medium to induce *PCH2* expression and a second sample of cells 3 h later (13 h post meiotic induction).

### Preparation of chromosome spreads from harvested pachytene cells

Chromosome spreads were prepared essentially as described in^37^. Harvested cells were spheroplasted to remove their cell wall before resuspending in MES wash (1 M sorbitol, 0.1 M MES, 1 mM EDTA, 0.5 mM MgCl_2_ pH 6.5). Cells were then lysed with 1% Lipsol detergent, before spreading their contents onto a glass microscope slide and adding a fixative (3% w/v paraformaldehyde, 3.4% w/v sucrose).

### Immunostaining spread pachytene chromosomes

Immunofluorescent labeling was performed as previously described^38^. Slides containing spread, fixed pachytene chromosomes, were incubated at room temperature in TBS buffer (25 mM Tris-Cl, pH 8, 136 mM NaCl, 3 mM KCl) then blocked with TBS buffer-1% w/v Bovine serum albumin (BSA). Primary antibodies (mouse monoclonal anti-Myc for Zip3 (Santa Cruz Biotechnology), goat polyclonal anti-Zip1 (Santa Cruz Biotechnology) rabbit polyclonal anti-Hop1 (generous gift from Franz Klein), or rat polyclonal anti-Zip2 (custom antibody, Abclonal) were diluted appropriately in TBS-1% BSA.

For *ndt80*Δ experiments involving labelling of Zip3, Hop1 and Zip1, secondary antibodies were donkey anti-mouse, anti-rabbit, and anti-goat IgG labeled with Alexa555 (for Zip3), Alexa488 (for Hop1), and Alexa647 (for Zip1) (Invitrogen Molecular Probes) and diluted appropriately in TBS-1% BSA. For *NDT80* experiments involving labelling of Zip2, Hop1 and Zip1, secondary antibodies were donkey anti-rat, anti-rabbit, anti-goat labeled with Alexa488 (for Zip2), Alexa555 (for Hop1), and Alexa647 (for Zip1). Chromosomal DNA was stained with DAPI (1 μg/ml) to permit detection/localization of pachytene chromosomes without the risk of bleaching the immunofluorescence signals of interest. Finally, the labelled chromosomes were mounted in Prolong Gold (Invitrogen Molecular Probes).

### Image Acquisition

Micrographs were obtained by widefield fluorescence microscopy using a Zeiss Axioplan 2ie MOT microscope, with an attached plan-apochromat 100x magnification, 1.4 numerical aperture, oil immersion objective (Zeiss), and a Hamamatsu ImageEM EM-CCD camera (model C9100-13; effective pixel size, 67 nm). The microscope was controlled by Metamorph (Molecular Devices) software.

### Obtaining signal intensity traces of spreads pachytene chromosomes

Signal intensity profiles of pachytene chromosomes were extracted from acquired images using open-source software FIJI^39^. Isolated chromosomes were traced along their mid-line using the segmented line selection tool with a 3-pixel width (Fig. 1B middle). Intensity values were extracted using the Plot Profile function and saved as .csv files for downstream analysis. Images were not pre-processed prior to measurement.

### Processing and Analysis of signal intensity profile

Signal intensity profiles acquired from ImageJ were imported into MATLAB using custom MATLAB function formatTriadData for downstream processing and analysis.

### Frequency Analysis

Fourier coefficients and (one-sided) Power Spectral Densities (PSDs) were calculated using custom MATLAB function triadFrequencyDomain which utilizes MATLAB’s fft function at its core. For each PSD, the two peaks with the highest power were determined using MATLAB’s findpeaks function. Distributions of high-powered PSD peaks were fit to three component Gaussian mixture models using MATLAB’s fitgmdist function. Custom MATLAB function triadFilterPlotProfileUsingFFT was used to create ‘reconstructed intensity profiles’ corresponding to the focal and domain components of each signal. Briefly, following conversion of signals into the frequency domain using Fast Fourier Transform, the power of frequencies either above (for focal component) or below (for domainal component) the stated thresholds were set to zero. Signals were then converted back into the spatial domain by inverse FFT utilizing MATLAB’s ifft function.

### Detection and downstream analysis of Focal and Domainal Peaks

Focal and domainal peaks (local maxima) were detected using default settings of the findpeaks function of MATLAB. To minimize noise detection, the smooth function of MATLAB was utilized to smooth focal signals with a moving average filter of 3 pixels. Distances between adjacent focal/domain peaks were calculated using custom MATLAB function getCOSpacing. For focal peaks, distributions were fit to single component gamma distributions using the gampdf function of MATLAB. For domain peaks, distributions were fit to two component gamma mixture models using function GMMestimator^40^. Custom MATLAB function getDomainalToFocalPeakDistances was used to calculated distances between domainal peaks and their adjacent focal peaks. Custom MATLAB function getDistanceToClosestPeak was used to calculate (closest) distances between triad components. Custom MATLAB function getNullModelIndependentPhasing was used to simulate expected results for independent phasing. Briefly, for each distance between adjacent (focal or domain) peak of the same protein (e.g. Zip3-Zip3), a number was calculated at random from a uniform distribution ranging from 0 to half that distance using the rand function of MATLAB. This was repeated 100 times. Coefficient of coincidence values were calculated using custom MATLAB function getCoC as previously described^15^. 30 intervals were used for analysis of focal peaks and 15 intervals were used for analysis of domainal peaks.

### Analysis of Genetic Crossovers

Published genetic crossover positions for yeast chromosome IV and XV was converted from base pairs DNA into μm chromosome length assuming 324 bp/nm. This conversion was based on the known length of these chromosomes (1.53 Mb and 1.05 Mb respectively) and measured average pachytene lengths (4.8 μm and 3.2 μm^16^), which give conversion factors of 319 nm/bp and 328 nm/bp respectively. 324 bp/nm is the average of these two values.

### Crossover Patterning Simulations

Crossover patterning was simulated using Beam-Film model software package MADpatterns v1.1^15, 41^ with additional custom MATLAB function sortMinorityCO_precSensitiv_retained as described below. The same parameters, based on those previously used to simulate crossover patterning on wild type yeast chromosome XV^16^, were used for all simulations except were specified. The precise workflow, including all parameters used, can be found in the custom MATLAB scripts noted below.

‘Simulation 1’ was carried out using custom MATLAB script twoTieredCrossoverPatterning_sim1. Briefly, an array of precursors (stress sensitive flaws) with associated (stress) sensitivity values was first created. Canonical crossover patterning was then simulated as previously described^15,16^ by application of a crossover designation force (Stress/’Smax’), that is counteracted by an inhibitory interference signal (stress relief and redistribution) that emanates from crossover-designated precursors (stress-sensitive flaws that converted to cracks) over a user-defined distance (‘L’) of 0.32 μm^16^. The final set of crossover-designated precursors at this stage were classified as canonical crossovers.

We propose that the second round of beam-film patterning utilizes the stress landscape that remains after the end of the first round of patterning (Fig. 6A). To mimic this, we first removed canonical designated precursors from the original precursor array. This allows us to determine the outcome when stress redistribution propagates past pre-existing cracks (i.e. via the beam/substrate) as implicit in our model. We then ran a second round of beam-film patterning on this new population of precursor arrays but with a reduced Smax to simulate the reduction of stress that occurs due to the first round of patterning (Fig. 6A); a compensatory increase in precursor sensitivity to re-engage the patterning process (Fig. 6A); and an increase in the distance over which stress redistribution propagates from 0.32 μm to 0.64 μm (Fig. 6A).

‘Simulation 2’ was carried out as previously described^15, 16^ using custom MATLAB script randomSprinklingMinorityCrossovers_sim. Briefly, canonical crossovers were first simulated as described above for ‘Simulation 1’. Next, each remaining precursor was converted into a minority crossover with a probability of 0.2.

### Signal Ratios

The signal intensity profiles of each molecule were normalized using custom MATLAB function normalizeSignalsBySTD which first subtracts the mean intensity of the profile, and then divides by the standard deviation of the profile. Then, for each chromosome, the ratios of Zip3/Hop1, Zip3/Zip1, and Hop1/Zip1 were calculated using custom MATLAB function getTriadSignalRatios. This function first adds the arbitrary value of 100 to each of the three signals (Zip3, Hop1 and Zip1), before calculating each ratio. Addition of an arbitrary value was used to avoid issues created by the presence of both positive and negative values.

### Population Average Signal Intensity Profiles

Population average signal intensity profiles of each molecule were obtained using custom MATLAB function getPopnAverageSignal. Briefly, each measured chromosome was first normalized to unit length by dividing each pixel position (in micrometers) by the chromosome length (in micrometers). To permit averaging, chromosomes must have a signal intensity measurement at the same normalized chromosome position as one another. This was achieved by fitting spline fitting followed by interpolation.

## Resource Availability

### Materials Availability

Yeast strains and antibodies are stored in the Kleckner Laboratory and can be requested from the lead contact.

### Data Availability

Primary image files and signal intensity profiles of traced chromosomes have been uploaded to Harvard Dataverse (https://dataverse.harvard.edu) and will be publicly available with an assigned DOI^42^ upon acceptance.

### Code Availability

Analysis code is publicly available from Github (https://github.com/mwhite4/multiscaleCrossoverPatterning). It will be published and assigned a citable DOI upon acceptance.

## Acknowledgements

We thank Kathleen Fleming and Fengfeng Zhuang for preliminary experiments; John Hutchinson for advice on stress-promoted patterning; Alison Grinthal for helpful discussions; Maria Mukhina for help with Fourier analysis; Liangran Zhang for yeast strains, and Franz Klein and Nancy Hollingsworth for anti-Hop1 antibodies. This work was funded by National Institute of Health grant R35GM136322 (N.E.K.) and Human Frontiers Science Program long-term fellowship LT000927/2013 (M.A.W.).

## Author contributions

Conceptualization, M.A.W., B.W., N.E.K.; Methodology, M.A.W., B.W., N.E.K.; Software, M.A.W.; Formal Analysis, M.A.W.; Investigation, B.W., L.C., G.L.; Resources, B.W., N.E.K.; Writing – Original Draft M.A.W. and N.E.K.; Writing – Review and Editing, M.A.W., B.W., G.L., N.E.K.; Visualization, M.A.W.; Supervision, N.E.K.; Funding Acquisition, N.E.K. and M.A.W.

---

**Supplementary Fig. 1.**
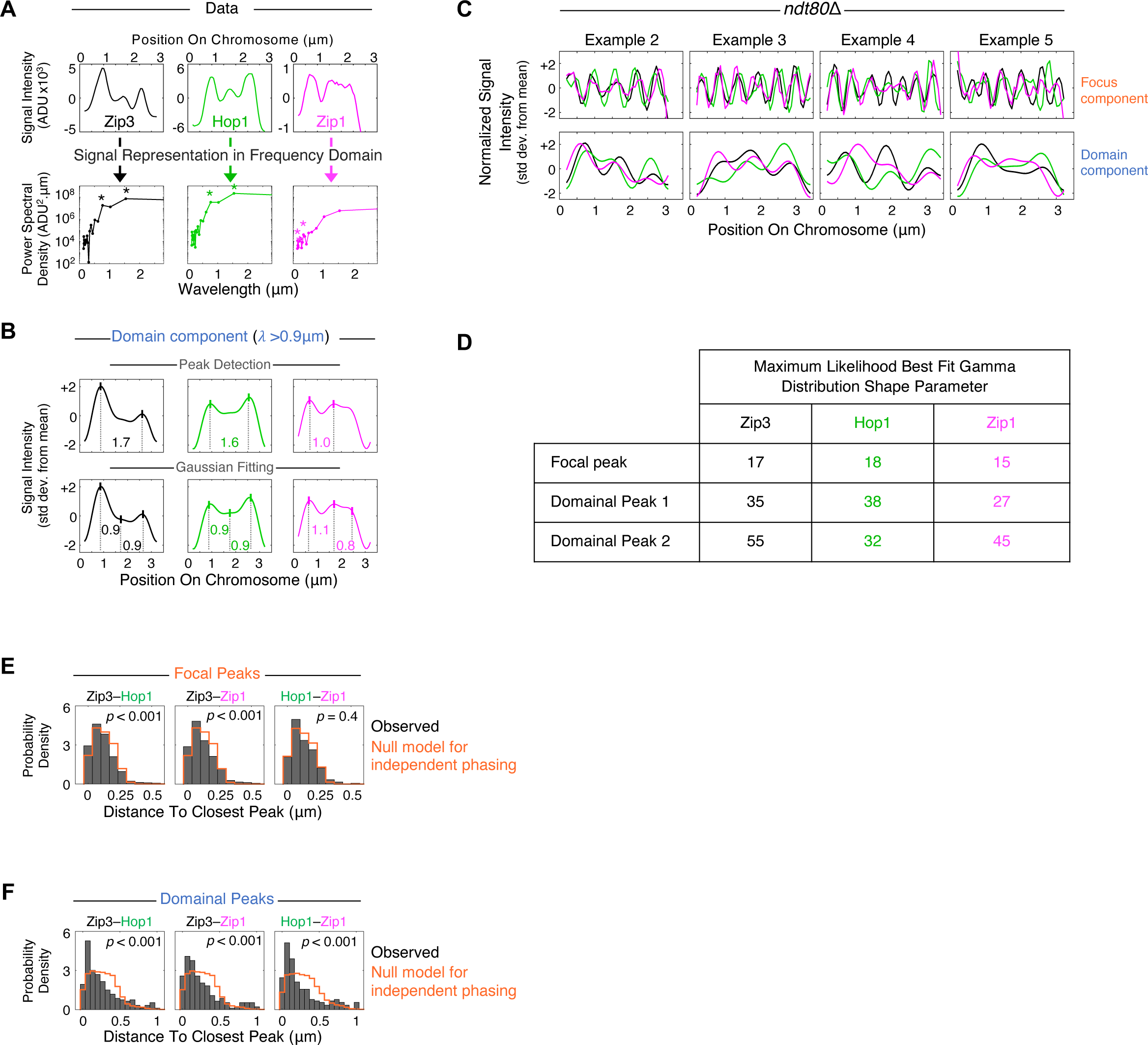
Frequency analysis reveals two component intensity periodicities corresponding to triads of focal and domainal signals, extended: part 1. *Related to* Fig. 2. (**A**) Intensity profiles and corresponding power spectral density plots of an example chromosome from a pachytene arrested *ndt80*Δ cell with high powered, long periodicity (1.54 μm) Zip3 and Hop1 signal components. Asterisks indicate two most prominent PSD peaks. ADU, analog digital units. (**B**) For this chromosome, the domain component of its signals has ‘peaks’ (vertical lines) that are not detected by standard peak detection (due their nature as ‘humps’) but are detected by Gaussian fitting. Distance between adjacent detected peaks indicated (μm). Std dev, standard deviations. (**C**) Normalized intensity profiles of focus and domain component of four additional example chromosomes. Std dev, standard deviations. (**D**) Shape parameters of maximum likelihood gamma distributions fit to distances between adjacent focal and domainal peaks (Fig. 2E, F). (**E** and **F**) Comparison of distributions of distances between each Zip3 peak and its closest Hop1 peak, with analogous distributions for Zip3-Zip1 and Hop1-Zip1 (‘Observed’; Fig. 2G, H) and null model for independent phasing of the two molecules (*see materials and methods*), with result of Wilcoxon rank sum test (*p*) indicated.

**Supplementary Fig. 2.**
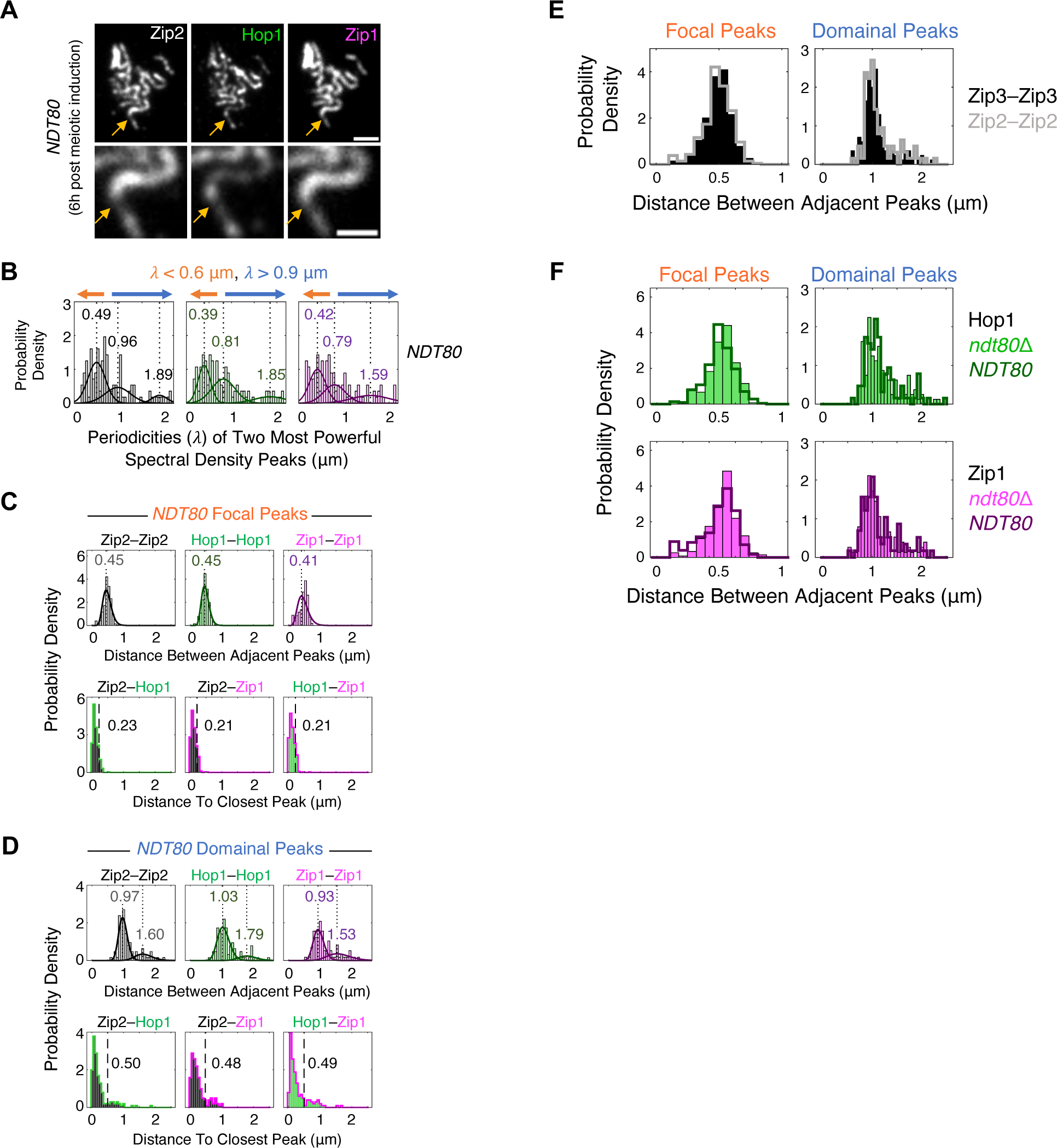
Frequency analysis reveals two component intensity periodicities corresponding to triads of focal and domainal signals, extended: part 2. *Related to* Fig. 2. (**A**) *Top*, immunostained spread chromosomes from a wild type (*NDT80*) pachytene cell. Arrows indicate example chromosome. Scale bar, 2 μm. *Bottom*, enlarged example chromosome. Scale bar, 1 μm. (**B**) Distributions of the two most powerful PSD peaks across all measured chromosomes (n = 56 chromosomes), with best fit 3-component Gaussian mixture models (solid lines) and their corresponding means (dashed lines). Orange and blue arrows indicate thresholds used for reconstruction of focal and domainal signals. (**C** and **D**) *Top*, distributions of distances between adjacent focal and domainal peaks, with best fit Gaussian distributions (solid lines) and their corresponding modes (dashed lines). *Bottom*, distributions of distances between each Zip2 peak and its closest Hop1 peak and analogous distributions for Zip2-Zip1 and Hop1-Zip1, for focal and domainal signal components. Dashed lines and corresponding values indicate expected medians if the two molecules were independently phased. (**E**) Comparison of distributions of distances between adjacent Zip3 and Zip2 focal and domainal peaks. (**F**) Comparison of distributions of distances between adjacent Hop1 and Zip1 focal and domainal peaks along chromosomes isolated from pachytene *NDT80* and pachytene arrested *ndt80*Δ cells.

**Supplementary Fig. 3.**
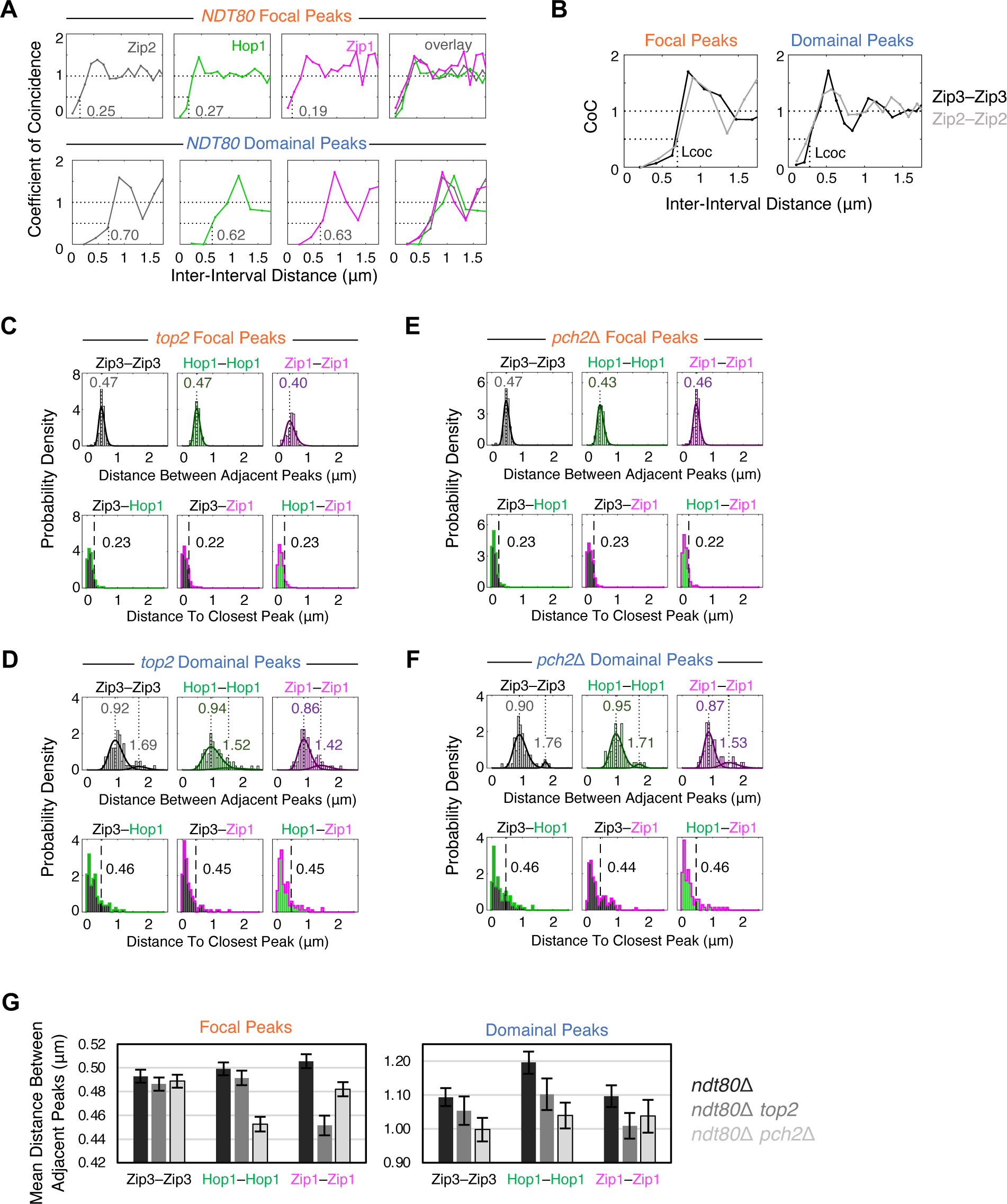
Crossover interference governs the patterns of both focal and domain triads, extended. *Related to* Fig. 3. (**A**) Coefficient of coincidence (CoC) curves for focal and domainal peaks of pachytene chromosomes isolated from a wild type strain in the absence of pachytene arrest (*NDT80*), with corresponding L_coc_ values. (**B**) comparison of CoC curves for Zip3 and Zip2 focal and domainal peaks. (**C** and **D**) *Top*, distributions of distances between adjacent focal and domainal peaks detected on chromosomes isolated from *top2* cells, with best fit Gaussian distributions (solid lines) and their corresponding modes (dashed lines) indicated. *Bottom*, distributions of distances between each Zip3 peak and its closest Hop1 peak detected on *top2* chromosomes, with analogous distributions for Zip3-Zip1 and Hop1-Zip1. Dashed lines and corresponding values indicate expected medians if the two molecules were independently phased. (**E** and **F**) as (**C** and **D**), except for focal and domainal peaks detected on chromosomes isolated from *pch2*Δ cells. (**G**) Mean distances between adjacent focal and domainal peaks for *ndt80*Δ, *ndt80*Δ *top2* and *ndt80*Δ *pch2*Δ cells. Error bars show standard error of the mean; n = 81, 60 and 52 for *ndt80*Δ, *ndt80*Δ *top2* and *ndt80*Δ *pch2*Δ cells respectively.

**Supplementary Fig. 4.**
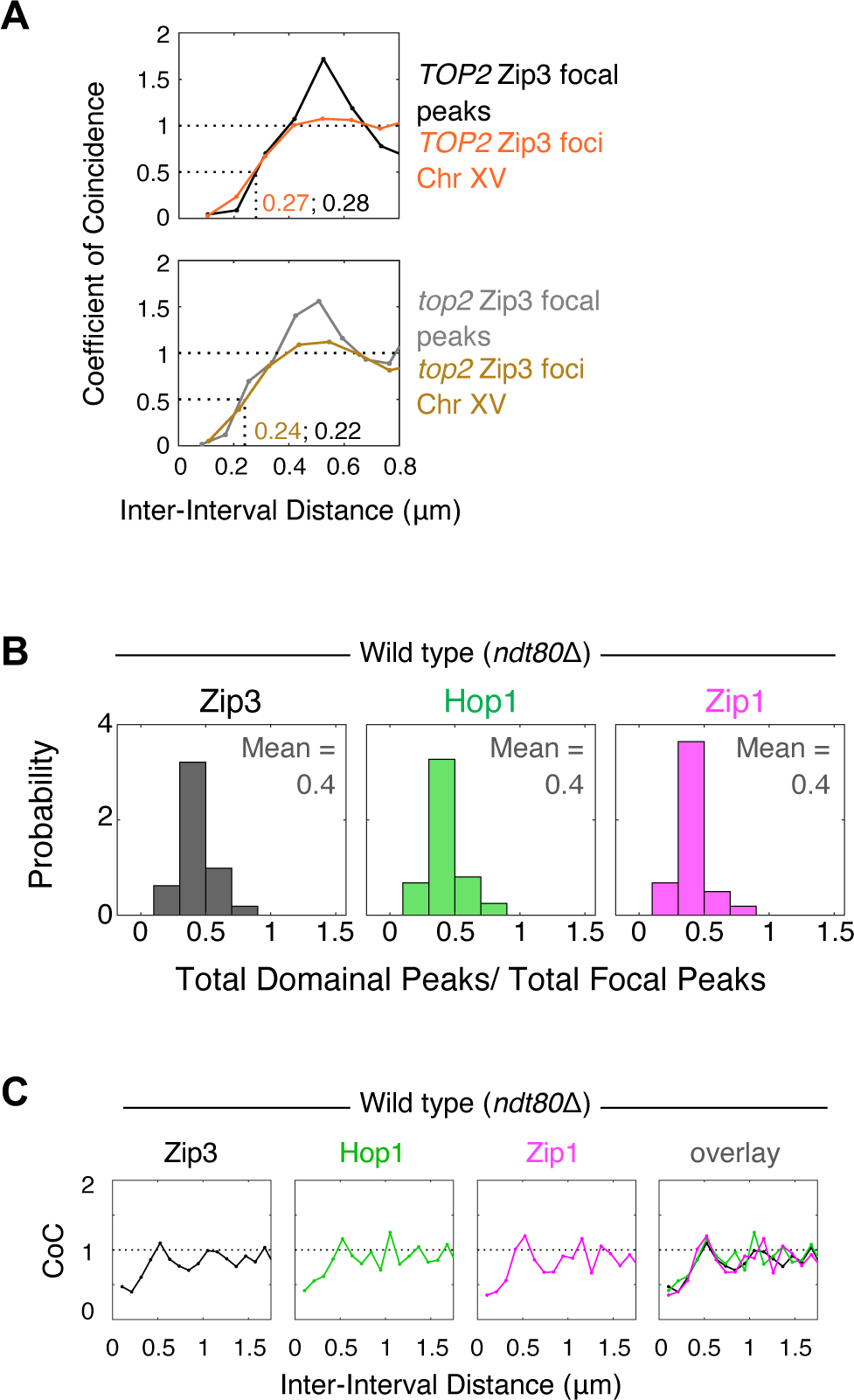
Focal triads correspond to canonical crossovers and domain triads correspond to minority crossovers, extended. *Related to* Fig. 4. (**A**) *top2* conditions confer the same reduction in L_CoC_, relative to wild type *TOP2* conditions for focal triad peaks (this work) as previously shown by cytological analysis of Zip3 foci^1^. ‘Chr’, chromosome. (**B**) Distribution of ratios of total domainal to total focal peaks in pachytene arrested *ndt80*Δ cells. (**C**) CoC curves for all peaks (focal and domanial) along measured chromosomes isolated from pachytene arrested *ndt80*Δ cells.

**Supplementary Fig. 5.**
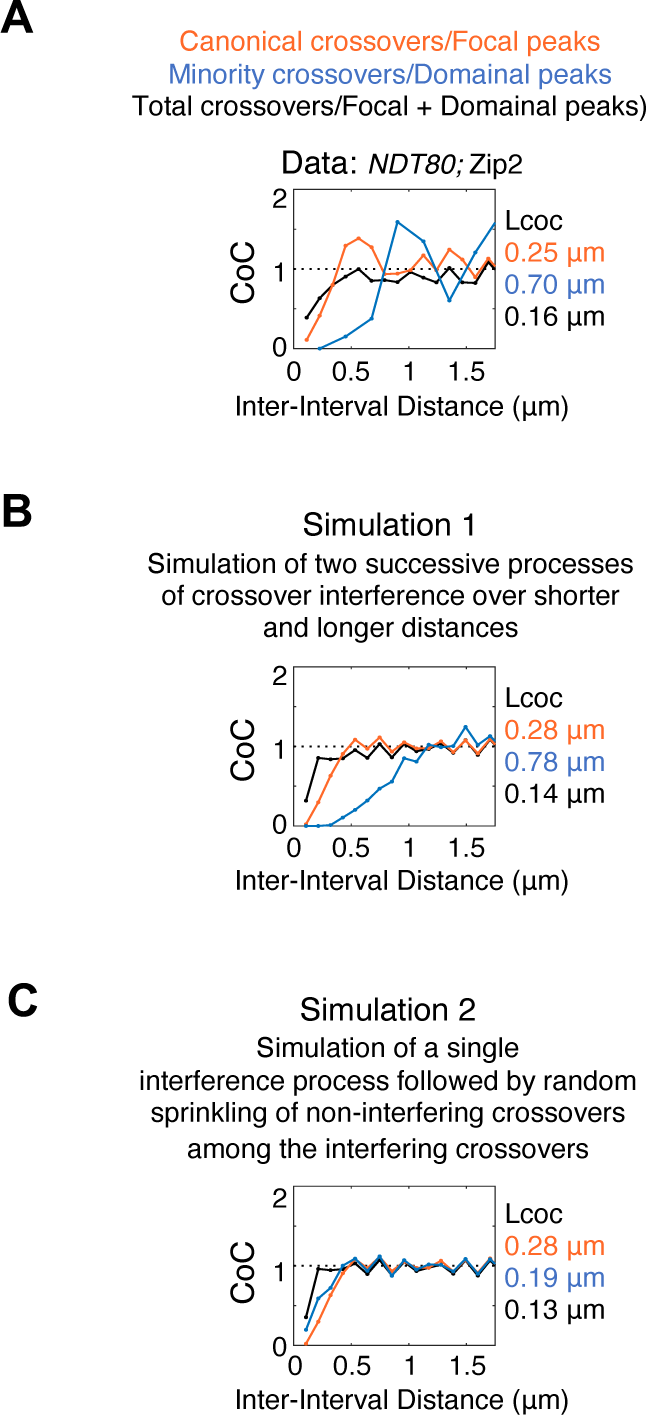
Simulations of crossover patterning. *Related to* Fig. 6. (**A** and **B**) CoC curves for triads as defined in this work (**A**) are well explained by simulations of two rounds of stress-promoted patterning as described in Fig. 6A (**B**). (**C**) Simulation of proposed model in which minority crossovers are non-interfering and are sprinkled randomly among canonical crossovers. Residual interference in the minority crossovers is due to even spacing of precursors (Supplementary note). Simulations performed as in^2^.

**Supplementary Fig. 6.**
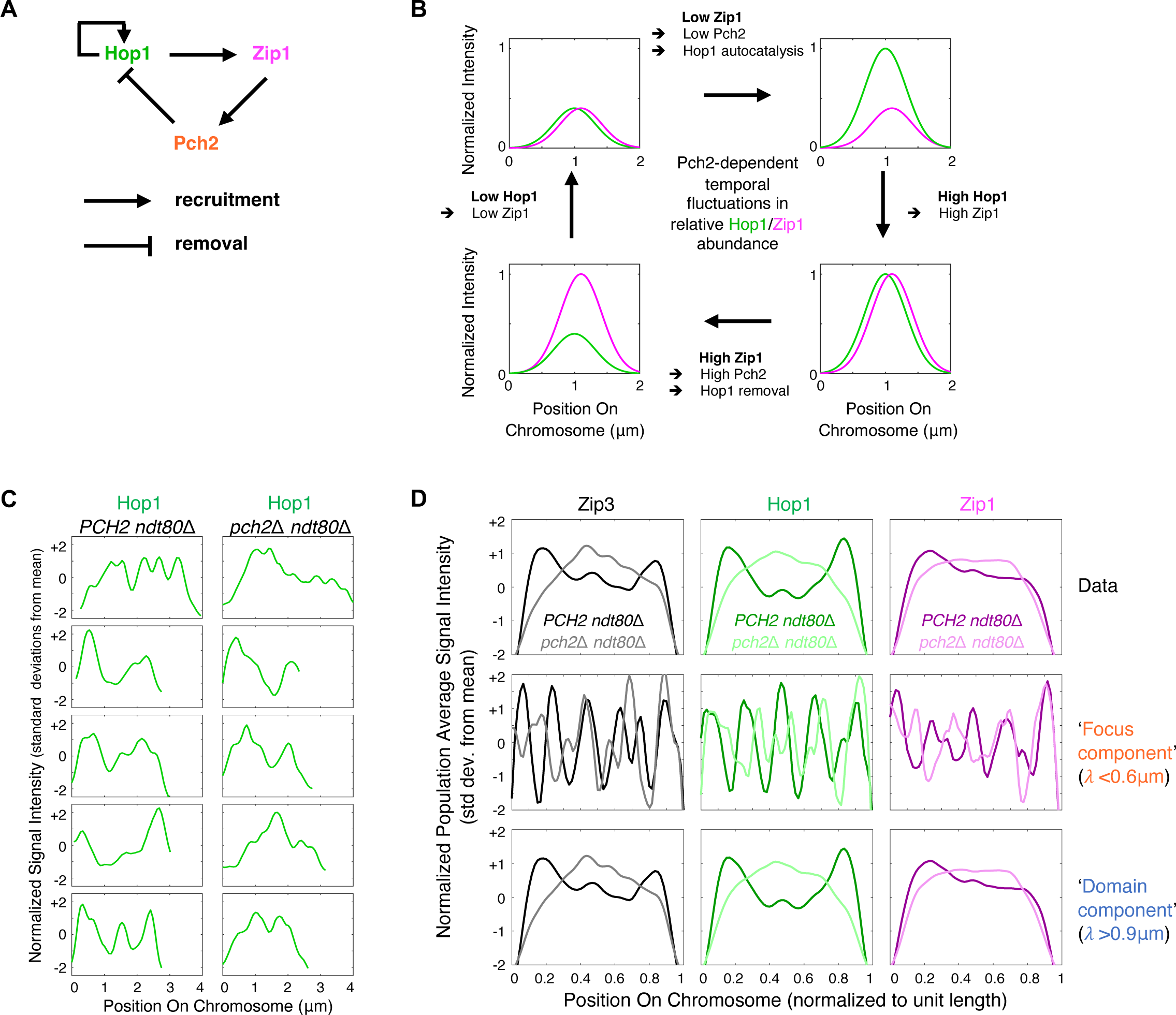
Pch2 uncouples Hop1 and Zip1 loading within triads, extended. *Related to* Fig. 5. (**A**) Model for Hop1, Zip1, Pch2 recruitment/removal interactions. (**B**) Schematic of temporal fluctuations in Hop1 and Zip1 abundances based on interaction model (**A**) and an assumption that Hop1-mediated Zip1 changes lag behind Hop1 changes. (**C**) Normalized Hop1 intensity profiles of five *PCH2 ndt80*Δ and five *pch2*Δ *ndt80*Δ example chromosomes. (**D**) Normalized intensity profiles of *PCH2 ndt80*Δ and *pch2*Δ *ndt80*Δ chromosomes averaged over all measured chromosomes (n = 81 and 52, respectively). Std dev, standard deviations.

## Supplementary Note

Average precursor spacing was derived from measured pachytene chromosomes lengths of wild-type yeast chromosomes III, XV and IV (1.2 μm, 3.2 μm and 4.9 μm respectively^3^); and their expected average number of total precursors based on Spo11 oligo counts and best fit crossover patterning simulations (6, 13, and 19 respectively^3^). The mean average predicted precursor spacing for these three chromosomes is 233 nm +/-29 nm (standard deviation). Simulations indicate that, consistent with cytological observations in other organisms^4, 5^, precursors are regularly spaced along the lengths of wild type yeast chromosomes^3^.

## Supplementary Discussion

It has often been stated that “Not all crossovers participate in interference”, with “some crossovers participating in interference and others neither contributing to nor responding to interference” (e.g. ^6^). This statement is not accurate, for the following reasons.

### Minority Crossovers in Wild Type Yeast Meiosis

As discussed in the text, it is clear that, in wild type yeast meiosis, the number of crossovers detected genetically or by DNA analysis of hybrid strains marked by sequence heterozygosities, is greater than the number of crossovers detected cytologically as Zip2/Zip3 foci. This disparity implies that, in wild type meiosis, there are two types of crossovers - the majority canonical set and a minority set that is detectable at the genetic/DNA level but is not detected cytologically as foci. However, the two types of crossovers cannot be distinguished from one another in such analyses, which only report total crossovers, without any indication as to which crossovers are of which type. Correspondingly, there is no way to determine directly from genetic/DNA data whether the minority subset does or does not exhibit interference.

It has been widely assumed that minority crossovers lack interference. However, this assumption has arisen largely from modeling (e.g. ^7^). Such modeling suggests that the total distribution of crossovers can be explained if there is a majority of crossovers that exhibit canonical interference (corresponding to those observed cytologically) and a minority of crossovers that are sprinkled “randomly” among those majority crossovers. The fundamental basis for the apparent success of such a simulation is that it can explain why, in the set of total crossovers, double crossovers along a single chromosome can be very closely spaced. One manifestation of this feature is the fact that the CoC curve for total crossovers does not fall to zero at small inter-interval distances (Fig. 4D). In the modeled scenario, these close double crossovers occur because a “non-interfering crossover” can occur very close to an “interfering crossover” or to another “non-interfering crossover”. However, we show above that the experimental CoC curve for total triads does not fall to zero at small inter-interval distances (Fig. 4D, E), even though interference for the minority crossovers is greater than that of canonical crossovers, rather than nonexistent. This feature is explained by the fact that crossovers/triads arising from the two tiers can occur at adjacent precursor sites, which are separated by ∼233 nm (above, ‘Precursor Arrays’) - the average distance between peaks in adjacent pairs of focal and domainal triads is the same as this distance, ∼270 nm (Fig. 4F).

The difference between the experimentally observed situation and the previously modeled situations is illustrated by mathematical simulations. Both simulations assume that all crossovers arise from the same set of precursors and that those precursors tend to be evenly spaced (above, ‘Precursor Arrays’). In the experimentally observed situation, CoC curves for total crossovers/triads, and for each of the two component sets of crossovers/triads can be simulated according to the two-stage scenario discussed in the text (Fig. 6A) and Supplementary Fig. 5B. In the previously modeled situation, the CoC curves for total crossovers are simulated by sprinkling minority crossovers/triads randomly among canonical crossovers/triads. The observed and previously modeled situations give quite similar CoC curves, with the appropriate diagnostic features seen in experimental data (Supplementary Fig. 5B versus 5C (black)). This is true even though, in the experimental data, minority crossovers/triads exhibit *more* interference than canonical crossovers/triads (Supplementary Fig. 5B, blue versus orange) whereas, in the previously modeled situation, minority crossovers have a lower level of interference than canonical crossovers (Supplementary Fig. 5C, blue versus orange). There is a residual level of interference of those “random” crossovers due to even spacing of precursor sites.

### Minority Crossovers in Wild Type Meiosis in Other Organisms

In *Arabidopsis thaliana,* maize, mouse and humans, wild type crossover patterns are well explained by the existence of two classes of crossovers that differ in interference behavior^7, 8, 9, 10, 11^. The minority crossovers in these cases have been suggested to lack interference for reasons related to those used previously for yeast and thus can also be explained by the two-tiered patterning described here. In tomato, the situation is more ambiguous. Comparison of late recombination nodules (thought to mark all crossovers) to Mlh1 foci (thought to specifically mark majority crossovers) resulted in the conclusion that ‘minority crossovers’ (i.e. late recombination nodules without a corresponding Mlh1 focus) lacked interference^12^. However, we note that this conclusion is based on the rare instances where chromosomes with more than one minority crossover per chromosome was identified (0.04% of all measured chromosomes). We also note that, in all the above cases, minority crossovers are rare (on average less than 1 per chromosome). Thus, a ‘typical’ scenario is occurrence of one or two majority crossovers and occasionally an extra minority crossover. Thus, direct measurement of interference between minority crossovers cannot be measured in these organisms. In principle, the interference distance for minority crossovers could be equal to, or longer than the chromosome length in these cases.

### Type II Crossovers in Mutants

The notion that the minority crossovers in wild type meiosis are “non-interfering” has also been inferred from the fact that absence of MutSγ (Msh4/Msh5) reduces the number of crossovers as compared to wild type meiosis and the crossovers that do form do not exhibit interference, and are resolved by non-canonical structure-specific nucleases (e.g. Mus81/Mms4 rather than MutLγ)^13^. It is thus sometimes assumed that a *mutSγ* mutation specifically subtracts canonical crossovers, thus leaving behind crossovers that are the same as the minority crossovers in wild type meiosis. If this were true, the minority crossovers in wild type meiosis would not exhibit interference. However, since we show that the minority crossovers in wild type meiosis DO exhibit interference, the “residual crossovers” in *mutSγ* mutants must not correspond to those wild type minority crossovers and must have a different origin, which explains why they do not exhibit interference. The *mutSγ* phenotype can be explained by proposition (suggested by N. Hunter and colleagues, personal communication) that MutSγ is required to arrest undifferentiated precursor recombination interactions so that they can undergo regulated differentiation into crossover and non-crossover types. In the absence of such arrest, progression will not be regulated and any crossovers that may occur, by whatever biochemical processes, will not exhibit interference.

The term Type II crossovers has also been applied to the crossovers that arise in RecQ-type helicase mutants. In such mutants, the number of crossovers is increased. Those crossovers do not exhibit interference and are resolved by non-canonical structure-specific nucleases^6^. In *Arabidopsis thaliana*, helicase-defective mutants exhibit very high levels of non-interfering crossovers; however, in such mutants, canonical crossovers still occur^14, 15^. The elevation in crossovers in helicase-defective mutants can therefore be attributed to aberrant progression of recombination interactions that would normally have become non-crossovers such that they often mature, instead, to crossover products (N. Hunter, personal communication). Such an effect is known to occur in such mutants by molecular studies of recombination in budding yeast mutants lacking RecQ helicase SGS1^16, 17^. In Arabidopsis, the number of non-crossovers is massively greater than the number of crossovers, so the result of this effect is a large increase in the total number of crossovers. And non-crossover-fated interactions, by their nature, are left over from the crossover designation/interference process and thus do not exhibit interference (except perhaps residual interference from even spacing of precursors). Thus, any crossovers arising from such interactions also will not exhibit interference. The same explanation explains the more modest elevation of crossovers seen in a yeast *sgs1* mutant^16, 17^. Thus, the crossovers that arise in *recQ* helicase mutants (which do not interfere) are also not related to the minority crossovers observed in wild type (which exhibit stronger interference than canonical crossovers).

